# TurboID-based proximity labeling discovers ABCF2 as an adhesion receptor for the zoonotic pathogen *Pasteurella multocida*

**DOI:** 10.1101/2024.12.03.626657

**Authors:** Fei Wang, Yuyao Shang, Menghan Chen, Zhihao Wang, Hanyuan Liu, Lin Hua, Wentao Li, Huanchun Chen, Qigai He, Bin Wu, Zhong Peng

## Abstract

*Pasteurella multocida* is a zoonotic pathogen that can cause fatal infections in both animals and humans. A significant number of putative adhesive factors have been predicted to contribute to the pathogenesis of *P. multocida*, but their interactive proteins on host cells remain unclear. In this study, we experimentally verified the roles of three previously proposed proteins (PlpE, PtfA, Hsf-2) in the adherence of *P. multocida*. Through turboID-based proximity labeling screening, we identified ATP-binding cassette sub-family F member 2 (ABCF2) as a host interactive protein for PlpE/PtfA/Hsf-2. Crucial amino acid residues in PlpE, PtfA, and/or Hsf-2 that are essential for interacting with ABCF2 included Asp-123 (PlpE), Lys-88 (PtfA), Asp-136 (PtfA), Ala-464 (Hsf-2), Glu-473 (Hsf-2), and Arg-489 (Hsf-2). Knocking down or knocking out ABCF2 significantly reduced the adherence and invasion of *P*. *multocida* to host cells, while overexpression of ABCF2 markedly increased these effects. However, ABCF2 did not contribute to the adherence of other bacterial species such as *Klebsiella pneumoniae* and *Bordetella bronchiseptica*. Additionally, we demonstrated that *P. multocida* infection upregulated the expression of host ABCF2 by activating the p38 MAPK and NF-κB signaling pathways. Furthermore, we showed that ABCF2 was involved in the *P. multocida*-induced p53-dependent apoptotic signaling pathway. To the best of our knowledge, this is the first identification of ABCF2 as a host factor contributing to the adherence of *P. multocida* and only the second report of ABCF2’s involvement in bacterial pathogenesis.

**Importance:** *P. multocida* can cause fatal infections in both animals and humans, yet the mechanisms related to its pathogenesis remain to be fully explored. In this study, we identified ABCF2 as a crucial host interactive protein for three adhesive proteins encoded by *P. multocida* and experimentally verified its role in the adherence and invasion of *P. multocida*. Furthermore, we elucidated how *P. multocida* modulates ABCF2 during its infection and revealed p53-dependent apoptosis as a downstream effect of ABCF2 during *P. multocida* infection. Given the absence of reports on ABCF2 contributing to the pathogenesis of *P. multocida* and only one previous report on the involvement of ABCF2 in bacterial pathogenesis prior to this study, this research could be valuable for comprehending the interactions between bacteria and hosts during bacterial infections.

## Introduction

Adhesion of pathogens to host cells or tissues is a universal prerequisite for bacteria to efficiently deploy their repertoire of virulence factors and cause infections (1). Consequently, bacteria have evolved various virulence factors to assist in binding to host cells during infection, including capsules, lipopolysaccharides (LPS), fimbriae, pili, and outer membrane proteins (OMPs) (2). On the surface of host cells, there exist numerous known or unknown receptors that can be recognized by bacterial adhesive proteins to establish interactions (3). Identifying important bacterial adhesive proteins and their corresponding host receptors is not only crucial for understanding bacterial pathogenesis but also aids in the development of anti-adhesion-based therapies for bacterial diseases (4).

*Pasteurella multocida*, a Gram-negative zoonotic bacterium that can cause fatal infections in humans and a wide range of animals, poses a significant threat to public health, leading to high morbidity, mortality, and economic losses in agriculture (5). Clinical strains of *P. multocida* are classified into five capsular genotypes (A, B, D, E, F) (6). Among these genotypes, types A and/or D are predominantly prevalent in various host species (7), and frequently lead to the respiratory symptoms (5). Indeed, *P. multocida* strains have been found to be able to disrupt the mammalian respiratory epithelial barrier (8). Similar to many other bacterial species, *P. multocida* strains are believed to encode multiple virulence factors involved in their adhesion to host cells, which may include capsules, fimbriae, other adhesins, and various OMPs (7). However, many of these adhesion proteins are identified through comparative genomic analysis and lack laboratory confirmation. Furthermore, limited knowledge exists regarding host receptors that may interact with these adhesion proteins.

Recently, TurboID-based proximity labeling has emerged as a novel approach for studying the spatial and interaction characteristics of proteins in living cells, enabling rapid and non-toxic proximity labeling within just 10 minutes (9). This innovative tool has been instrumental in identifying host effector proteins responding to pathogen virulence factors, thereby enhancing our understanding of the pathogenesis of various pathogens, including viruses, bacteria, and parasites (10–12). In this study, we employed TurboID-based proximity labeling to screen for host receptors that potentially play crucial roles in the adhesion and pathogenesis of *P. multocida*. Notably, we discovered ATP-binding cassette sub-family F member 2 (ABCF2) as an adhesion receptor for *P. multocida* on mammalian respiratory epithelial cells. For the first time, we found ABCF2 involves in *P. multocida* induced p53-dependent apoptotic signaling pathway.

## Results

### ABCF2 is identified as a host factor to interact with different *P. multocida* adhesion proteins and plays an important role in mediating *P. multocida* adhesion

Previous genomic analyses have identified the type 4 fimbriae subunit PtfA, surface fibril Hsf-2, and *Pasteurella* lipoprotein E (PlpE) as adhesion proteins in *P. multocida* (7, 13, 14). To experimentally validate these predictions, we cloned the genes encoding these proteins into *E. coli* using the pET-28a(+) vector and conducted cell adhesion assays. The results demonstrated that *E. coli* strains expressing the aforementioned *P. multocida* adhesion proteins exhibited significantly enhanced capabilities in adhering to mammalian cells compared to strains carrying empty plasmids or control strains (Fig. 1A, B). Furthermore, overexpression of *ptfA*, *hsf-2*, and *plpE* in *P. multocida* also increased bacterial adhesion to various mammalian respiratory epithelial cells, including human lung epithelial cells (A549), swine trachea epithelial cells (NPTr), and human trachea epithelial cells (BEAS-2B) (Fig. 1C, D, E). To further support these findings, we deleted *plpE* from the *P. multocida* strain HuN001 (GenBank accession no. CP073238) and compared the cell adhesion and invasion capacities between the wild-type strains and the *plpE*-deleted strain (ΔPlpE). The results revealed that ΔPlpE exhibited significantly reduced cell adhesion and/or invasion capabilities compared to the wild-type strains (Fig. 1F, G, H, I, J). *In vivo* tests in mouse models also showed that ΔPlpE exhibited a significantly decreased capacity for adhering to and invading murine lungs compared to the wild type strain (Fig. 1K). These experiments provided experimental confirmation of the adhesion functions of PtfA, Hsf-2, and PlpE in *P. multocida*.

**Fig. 1.**
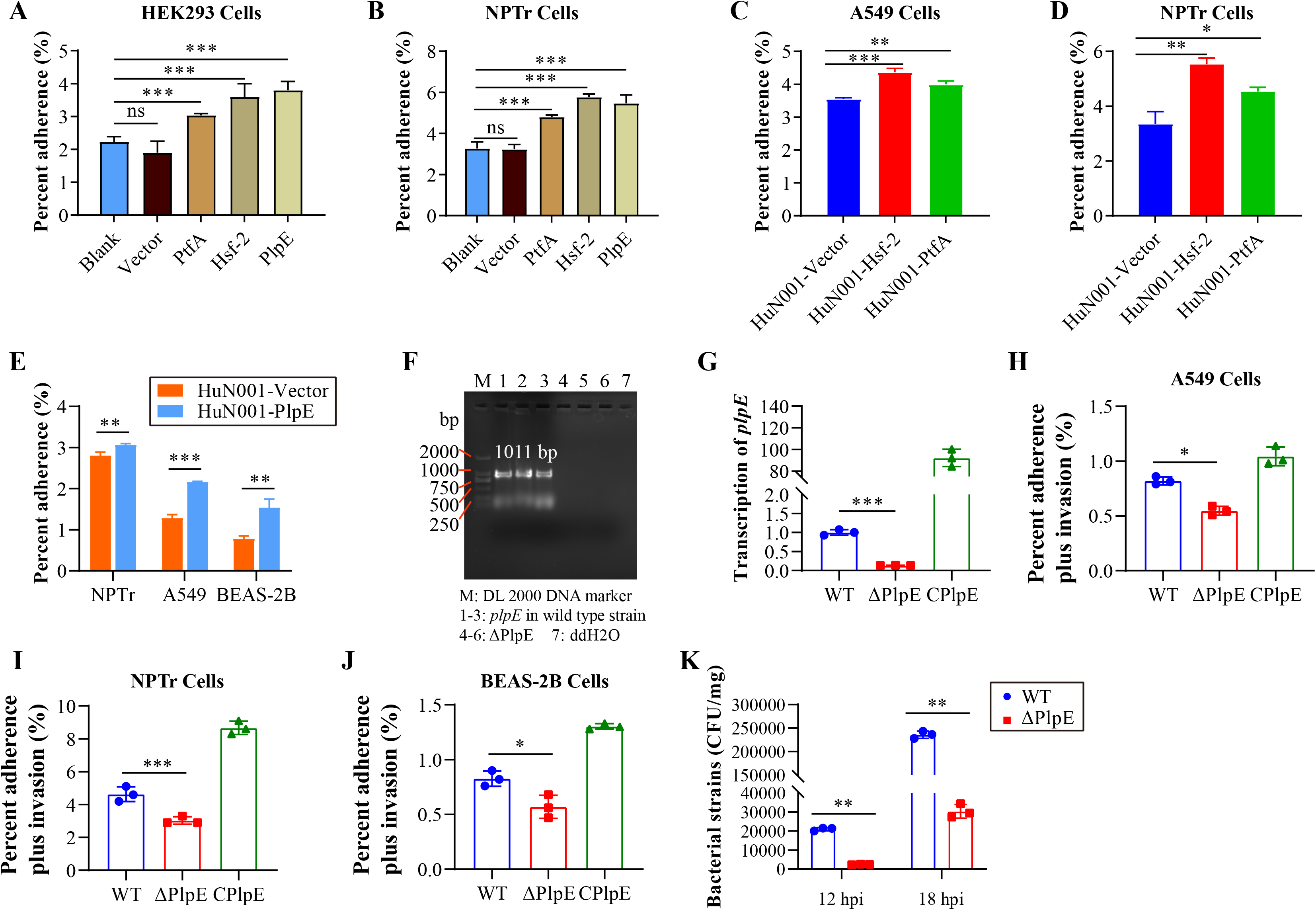
Experimental validation of different proteins contributing to the adherence of *Pasteurella multocida*. **(A∼E)** Column charts showing percent strains of different bacteria adhering to HEK-293T cells **(A)**, NPTr cells **(B, D, E)**, A549 cells **(C, E)**, and/or BEAS-2B cells **(E)**. In panels **A** and **B**, the X-axis refers to *E. coli* DH5α (Blank), DH5α harboring pET-28a(+) (Vector), DH5α harboring pET-28a(+) bearing *plpE* (PlpE), *ptfA* (PtfA), and/or *hsf-2* (Hsf-2) from *P. multocida*. In panels **C** and **D**, the X-axis refers to *P. multocida* HuN001 harboring pPBA1101 (HuN001-Vector), HuN001 harboring pPBA1101 bearing *ptfA* (HuN001-PtfA), and/or *hsf-2* (HuN001-Hsf-2). **(F)** PCR verification for the deletion of *plpE* from *P. multocida* HuN001. **(G)** A column chart showing the examination of transcriptional level of *plpE* in *P. multocida* wild type strain (WT), the *plpE*-deleted strain (ΔPlpE), and *plpE*-complementary strains (CPlpE). **(H∼J)** Column charts showing percent *P. multocida* strains adhering to A549 cells **(H)**, NPTr cells **(I)**, and BEAS-2B cells **(J)**. **(K)** A column chart showing the colony forming unit counts of *P. multocida* wild type strain (WT) and *plpE*-deleted strain (ΔPlpE) recovered from murine lungs at 12 hours post infection (hpi) and 18 hpi. In all column charts, data were presented as mean ± standard deviation (SD). The significance level was set at *P* > 0.05 (no significance [ns]), *P* < 0.05 (*), *P* < 0.01 (**), or *P* < 0.001 (***).

Subsequently, we utilized TurboID-based proximity labeling to screen for host receptors for *P. multocida* PlpE, PtfA, and/or Hsf-2, following a previously established protocol (15). Mass spectrometric analysis identified numerous host effector proteins potentially interacting with *P. multocida* PlpE, PtfA, and/or Hsf-2 in NPTr cells (Fig. 2A). Notably, ABCF2 was consistently present in the candidate lists for all three adhesion proteins with high scores (Fig. 2B), prompting its selection for further investigation. Indirect immunofluorescence (IIF) detection confirmed the expression of ABCF2 on the surface of NPTr and A549 cells (Fig. 2C). Further analyses using co-immunoprecipitation (Co-IP) and bio-layer interferometry (BLI) revealed clear interactions between ABCF2 and *P. multocida* PlpE, PtfA, and/or Hsf-2 (Fig. 2D, E). Additionally, colocalizations were observed between ABCF2 and PlpE (Fig. S1).

**Fig. 2.**
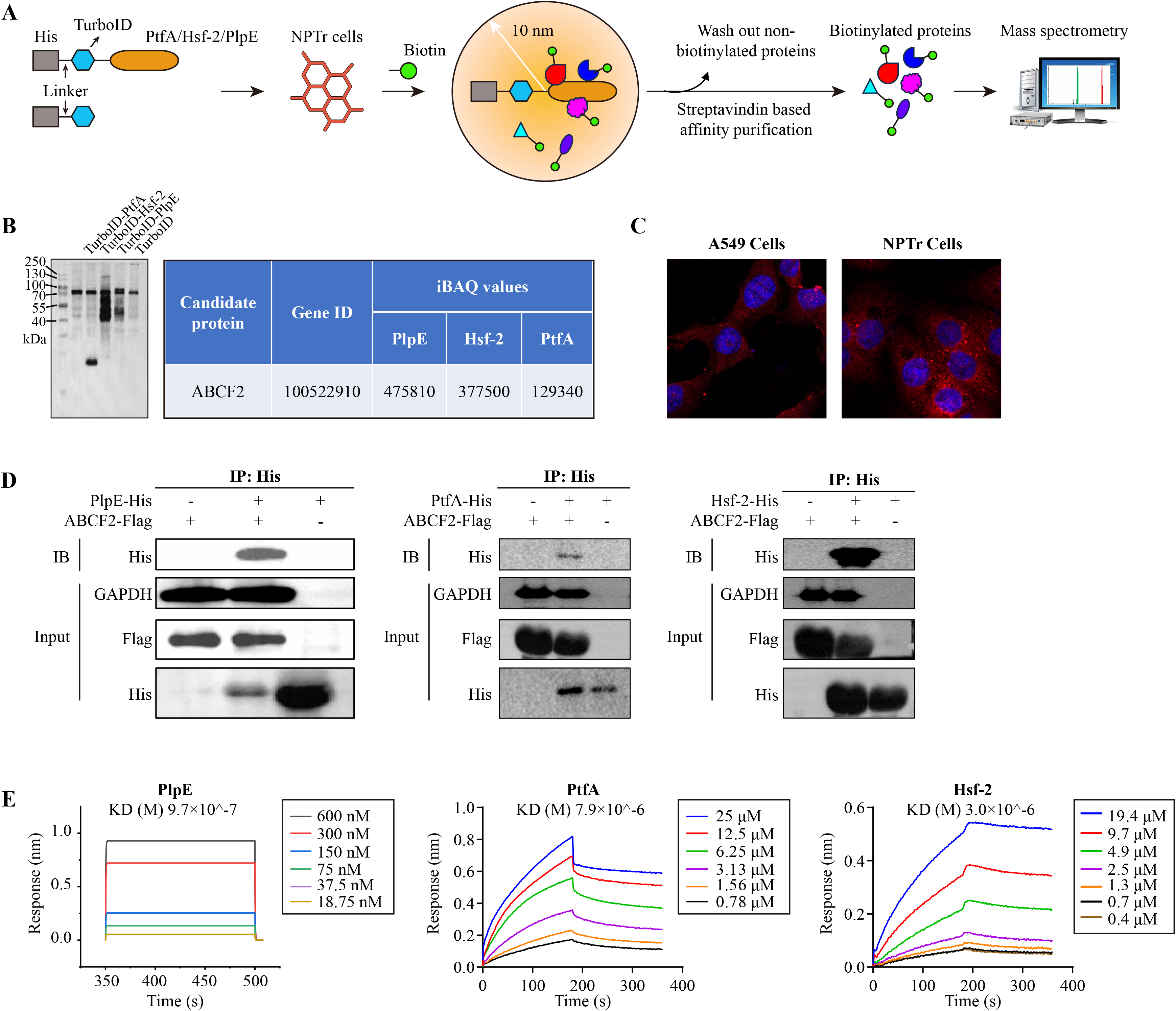
TurboID screening identified ABCF2 as a host protein interacting with different *Pasteurella multocida* adhesive proteins. (A) A diagram showing the workflow for TurboID screening. **(B)** An SDS-PAGE image showing the gel patterns of different recombinantly expressed proteins used for mass spectrometric analysis, and the iBAQ values of ABCF2 against different *P. multocida* adhesive proteins. **(C)** Validation of the expression of ABCF2 in A549 and NPTr cells by immunofluorescence assay. **(D)** Validation of the interactions between ABCF2 and different *P. multocida* adhesive proteins (PlpE, PtfA, and Hsf-2) by co-immunoprecipitation (Co-IP) assays. **(E)** Validation of the interactions between ABCF2 and different *P. multocida* adhesive proteins (PlpE, PtfA, and Hsf-2) by bio-layer interferometry (BLI) assays.

To evaluate the impact of ABCF2 on *P. multocida* adhesion, we overexpressed ABCF2 in NPTr and A549 cells and assessed bacterial adhesion using *P. multocida* strains of different genotypes. The results demonstrated a significant increase in the adhesion of various *P. multocida* strains to NPTr and A549 cells upon ABCF2 overexpression (Fig. 3A, B). Subsequently, we knocked down ABCF2 in NPTr cells, and cell adhesion assays indicated a significant decrease in the adhesion of different *P. multocida* strains to the cells (Fig. 3C, D). However, ABCF2 knockdown did not affect the adherence of other respiratory bacterial species, including *Bordetella bronchiseptica* and *Klebsiella pneumoniae*, to NPTr cells (Fig. S2). These findings emphasize the essential role of ABCF2 in mediating *P. multocida* adhesion to host cells.

**Fig. 3.**
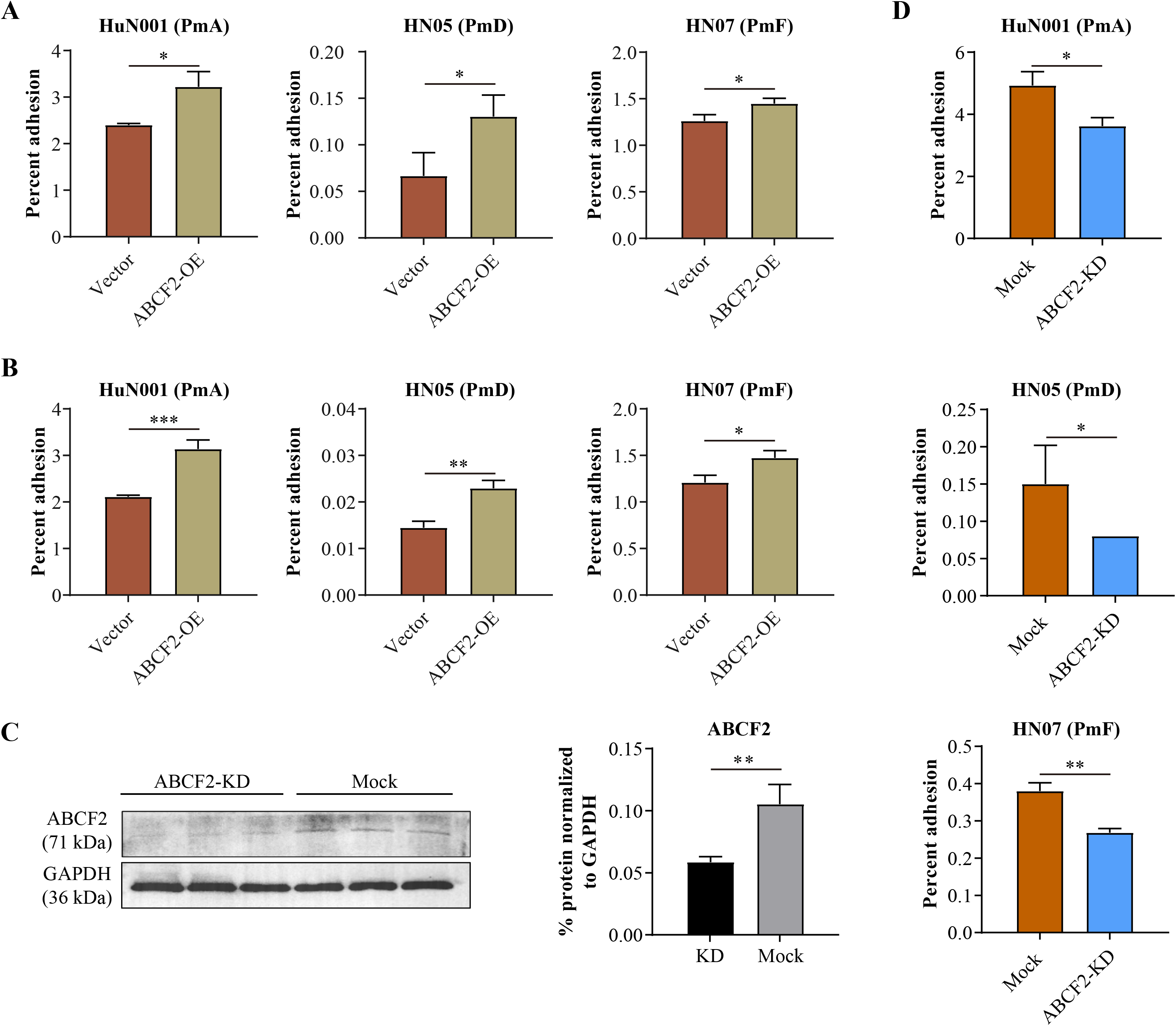
Validation of host ABCF2 contributing to *Pasteurella multocida* adherence. **(A, B)** Column charts showing percent *P. multocida* strains belonging to different serogroups adhering to NPTr cells with ABCF2 overexpressed **(A)**, and A549 cells with ABCF2 overexpressed **(B)**. In panel **A**, the X-axis refers to NPTr cells containing the vector plasmid (Vector) and overexpressed *abcf2* using a plasmid (ABCF2-OE). In panel **B**, the X-axis refers to A549 cells containing the vector plasmid (Vector) and overexpressed *abcf2* using a plasmid (ABCF2-OE). **(C)** Western-blotting results verifying the knocking down (KD) of ABCF2 in NPTr cells using a commercially synthesized siRNA. **(D)** showing percent *P. multocida* strains belonging to different serogroups adhering to NPTr wild type cells (mock) and/or ABCF2-knocking down cells (ABCF2-KD). In all column charts, data were presented as mean ± standard deviation (SD). The significance level was set at *P* < 0.05 (*), *P* < 0.01 (**), or *P* < 0.001 (***).

### Key amino acids for interacting with ABCF2 in *P. multocida* PlpE, PtfA, and Hsf-2

To further explore the interactions between ABCF2 and *P. multocida* adhesion proteins, we conducted molecular docking analysis and utilized AlphaFold3 (16) to predict the essential amino acids involved in the interaction with ABCF2 in PlpE, PtfA, and/or Hsf-2. This computational approach unveiled a set of key amino acids deemed crucial for the interaction between PlpE, PtfA, or Hsf-2 and ABCF2. These residues encompassed Asp-123, Ser-186, Glu-321, Thr-292, Asn-296, and Thr-298 in PlpE; Tyr-36, Tyr-39, Arg-56, Tyr-64, Asn-65, Gly-78, Arg-80, Lys-88, Ala-121, Gln-123, Thr-128, Val-129, and Asp-136 in PtfA; and Ala-464, Glu-473, Ala-478, Arg-489, and Ala-499 in Hsf-2 (Fig. 4A, B, C and Table S1). Subsequently, we generated and expressed a series of truncated proteins containing the identified amino acid residues, and evaluated their interactions with ABCF2 through Co-IP assays (Fig. 4D, E, F, G). The outcomes highlighted the significance of Asp-123 in PlpE, Lys-88 and Asp-136 in PtfA, as well as Ala-464, Glu-473, and Arg-489 in Hsf-2 for interacting with ABCF2. Further co-IP analyses using proteins containing mutated amino acid residues at these sites confirmed their critical roles in interacting with ABCF2 (Fig. 4H, I).

**Fig. 4.**
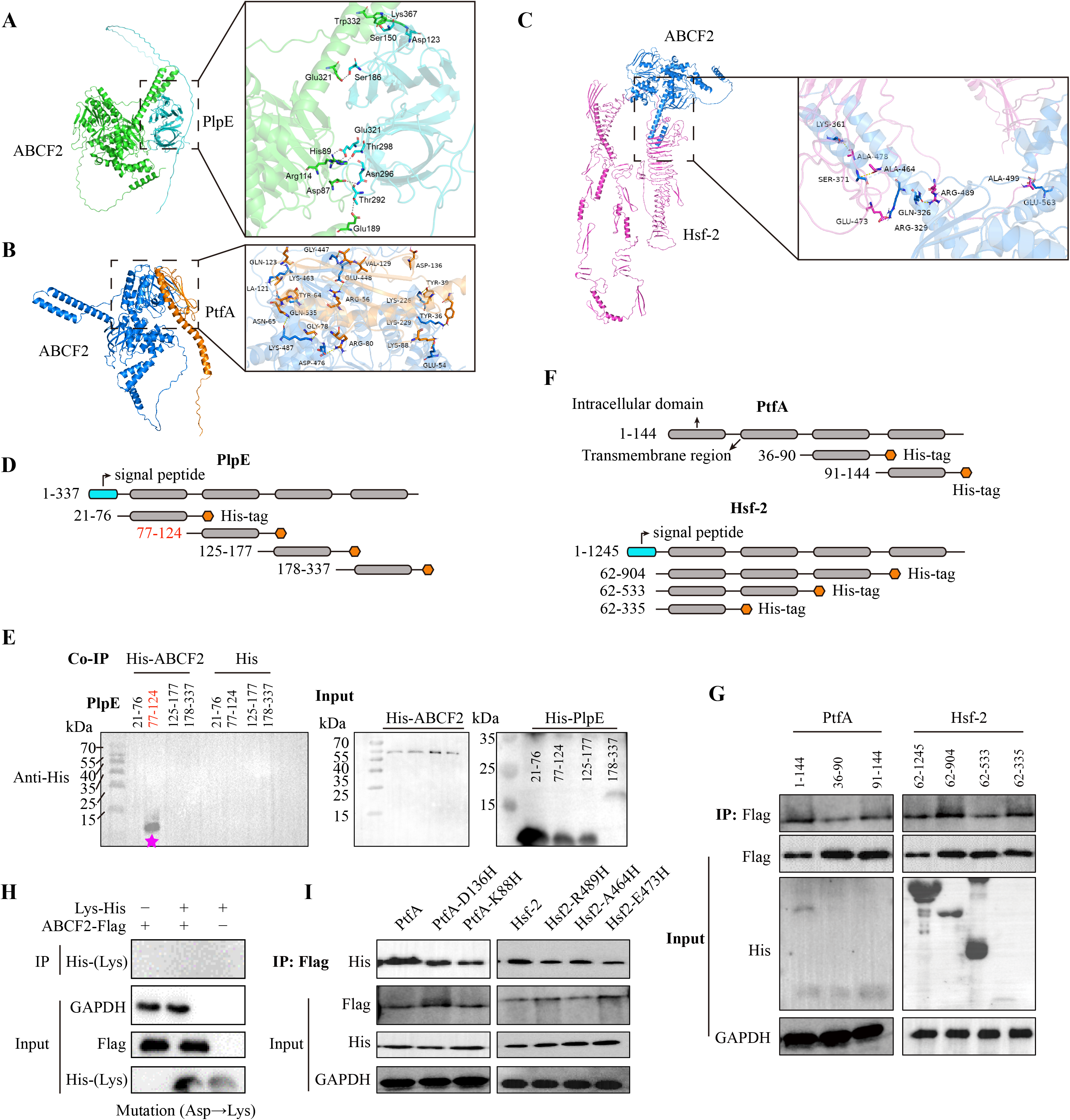
Identification and preliminary verification of amino acid residues crucial for interacting with ABCF2 in *Pasteurella multocida* adhesive proteins. (A∼C) Molecular docking between ABCF2 and *P. multocida* adhesive protein PlpE **(A)**, PtfA **(B)**, and Hsf-2 **(C)** using the structures predicted by AlphaFold3. Predicted amino acid residues crucial for interacting with ABCF2 in *P. multocida* adhesive proteins are shown in the boxes. **(D, F)** Strategies used for designing truncated proteins containing the identified amino acid residues by molecular docking analysis. **(E, G)** Validation of the interactions between ABCF2 and different truncated proteins generated from PlpE **(E)**, PtfA **(G)**, and Hsf-2 **(G)** by co-immunoprecipitation (Co-IP) assays. **(H, I)** Validation of the interactions between ABCF2 and different proteins containing mutated amino acid residues generated from PlpE **(H)**, PtfA **(I)**, and Hsf-2 **(I)** by co-immunoprecipitation (Co-IP) assays.

### *P. multocida* infection promotes ABCF2 expression through elevating the NF-κB and p38 MAPK signaling

We then evaluated the expression of ABCF2 following *P. multocida* inoculation. Western blot and qPCR analyses indicated that inoculation with *P. multocida* induced the expression of ABCF2 (Fig. 5A, B). However, knockout of *plpE* led to a significant decrease in ABCF2 expression (Fig. 5C, D). Given the reported regulation of ABCF2 by the nuclear factor-kappa B (NF-κB) signaling pathway in LPS-induced ulcerative colitis models (17), we explored the impact of NF-κB on ABCF2 regulation post *P. multocida* infection. The findings revealed a notable increase in P65 phosphorylation (P-P65) at 6 hpi following *P. multocida* infection, preceding the time point at which ABCF2 elevation was triggered by *P. multocida* (Fig. 5E, F). Significantly, treatment with the NF-κB inhibitor (BAY11-7082) markedly reduced the expression of ABCF2 induced by *P. multocida* infection (Fig. 5G, H), indicating the involvement of NF-κB in regulating ABCF2 expression post *P. multocida*infection.

**Fig. 5.**
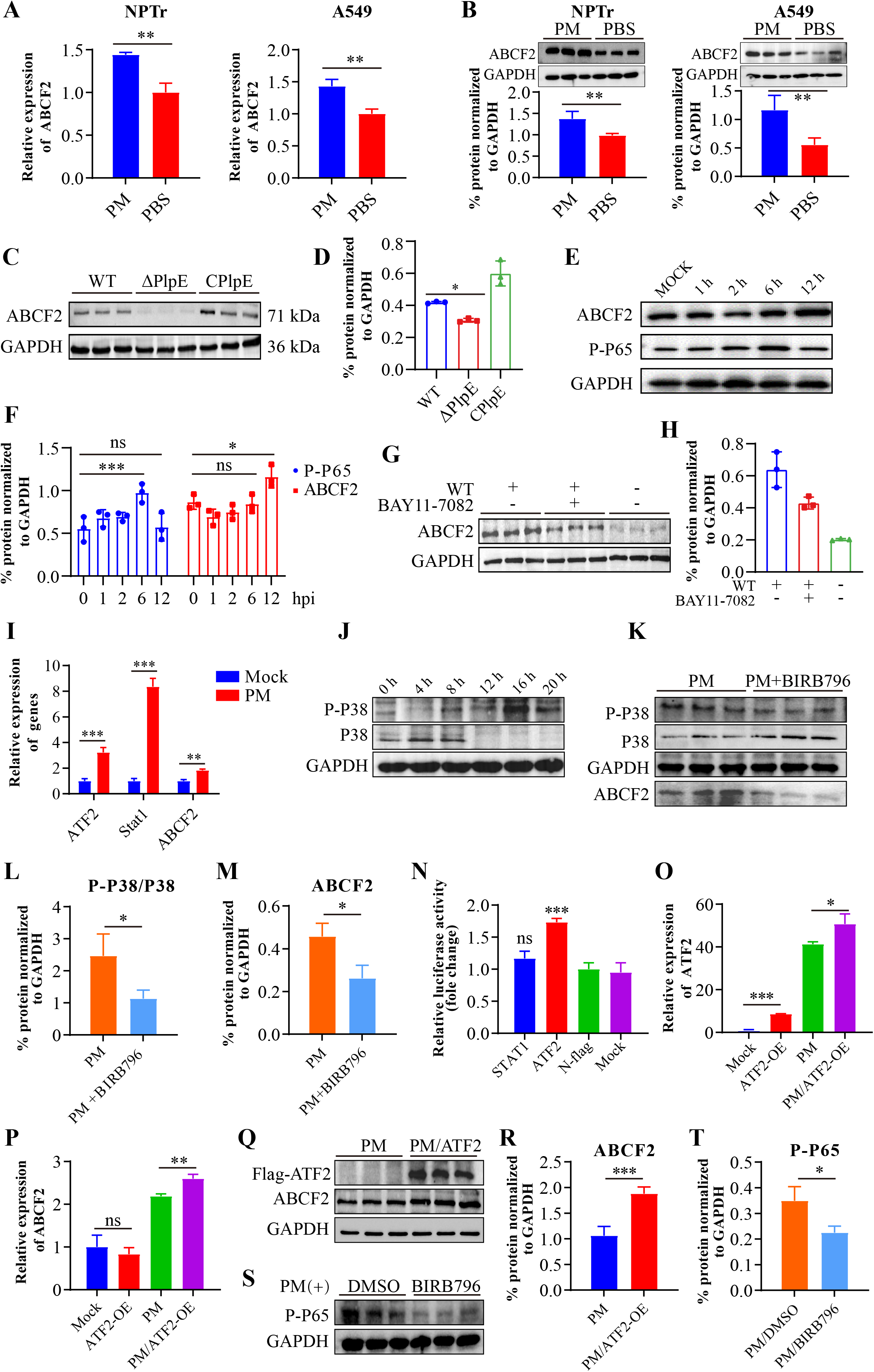
*Pasteurella multocida* infection promotes ABCF2 expression through elevating the NF-κB and p38 MAPK signaling. **(A, B)** *P. multocida* infection induced increased levels of ABCF2 in NPTr and A549 cells evaluated by qPCR **(A)** and western-blotting **(B)** assays. **(C)** Western-blotting results showing the expression of ABCF2 in NPTr cells induced by *P. multocida* wild type strain (WT), the *plpE*-deleted strain (ΔPlpE), and *plpE*-complementary strains (CPlpE). **(D)** Results from western-blotting assays quantified by the image J software. **(E)** Western-blotting results showing the expression of ABCF2 and P65 phosphorylation (P-P65) in NPTr cells at 0 (mock), 1, 2, 6, and 12 hours post *P. multocida* inoculation. **(F)** Results from western-blotting assays quantified by the image J software. **(G)** Western-blotting results showing the expression of ABCF2 and P65 phosphorylation (P-P65) in NPTr cells treated with (+) or without (-) the NF-κB inhibitor (BAY11-7082), followed by *P. multocida* inoculation. **(H)** Results from western-blotting assays quantified by the image J software. **(I)** qPCR detecting the transcriptional levels of *atf2*, *stat1*, and *abcf2* in NPTr cells at 12 hours post *P. multocida* inoculation. **(J)** Western-blotting results showing the P38 phosphorylation (P-P38) in NPTr cells at 0, 4, 8, 12, 16, and 20 hours post *P. multocida* inoculation. **(K)** Western-blotting results showing the expression of ABCF2 and P38 phosphorylation (P-P38) in NPTr cells treated with (+) or without (-) p38 inhibitor (BIRB796), followed by *P. multocida* inoculation. **(L)** P38 phosphorylation (P-P38) from western-blotting assays quantified by the image J software. **(M)** ABCF2 expression from western-blotting assays quantified by the image J software. **(N)** Dual luciferase assays demonstrating the regulation of *atf2* on the expression of *abcf2*. **(O)** qPCR assays verifying ATF2 overexpression (ATF2-OE) in NPTr cells with or without *P. multocida* infection. **(P)** qPCR assays showing ATF2 overexpression (ATF2-OE) contribute to ABCF2 expression after *P. multocida* infection. **(Q)** Western-blotting revealing ATF2 overexpression (ATF2-OE) contribute to ABCF2 expression after *P. multocida* infection. **(R)** ABCF2 expression from western-blotting assays quantified by the image J software. **(S)** Western-blotting showing inhibition of p38 MAPK signaling using the inhibitor BIRB796 decreases P65 phosphorylation (P-P65) after *P. multocida* infection. **(T)** The level of P65 phosphorylation (P-P65) from western-blotting assays quantified by the image J software. In all column charts, data were presented as mean ± standard deviation (SD). The significance level was set at *P* > 0.05 (no significance [ns]), *P* < 0.05 (*), *P* < 0.01 (**), or *P* < 0.001 (***).

Several previous studies have underscored the association between ABCF2 and cell apoptosis (18, 19). Considering the crucial role of p38 mitogen-activated protein kinase (MAPK) signaling in apoptosis regulation (20), we investigated whether p38 MAPK signaling is implicated in the regulation of ABCF2 post *P. multocida* infection.

The results demonstrated that *P. multocida* infection activated the p38 MAPK signaling pathway, while inhibition of p38 MAPK decreased the expression of ABCF2 induced by *P. multocida* (Fig. 5I, J, K, L, M). Our dual luciferase assay revealed that ATF2, a transcription factor activated by p38 MAPK, could bind to the promoter of ABCF2 (Fig. 5N). Additionally, *P. multocida* infection led to an increase in ATF2 expression, ultimately resulting in elevated ABCF2 expression (Fig. 5O, P, Q, R). However, overexpression of ATF2 alone did not stimulate the expression of ABCF2 (Fig. 5O, P, Q, R). These findings suggest that *P. multocida* infection enhances ABCF2 expression by activating the p38 MAPK signaling pathway. Moreover, p38 MAPK signaling has also been reported to play a role in regulating the NF-κB signaling (21). Subsequently, we investigated the relationship between p38 MAPK signaling and NF-κB signaling in *P. multocida*-induced ABCF2 expression. To do so, we inhibited p38 MAPK signaling using a commercially available inhibitor, BIRB796, and examined the level of P-P65 induced by *P. multocida* infection. The results showed that inhibition of p38 MAPK signaling significantly decreased the level of P-P65 after *P. multocida* infection (Fig. 5S, T), suggesting the involvement of p38 MAPK-regulated NF-κB signaling in this process. In summary, the aforementioned findings indicate that *P. multocida* infection promotes ABCF2 expression by upregulating the NF-κB and/or p38 MAPK/NF-κB signaling axis.

### ABCF2 involves in *P. multocida* induced p53-dependent apoptotic signaling pathway

Given the well-documented association between ABCF2 and apoptosis (18, 19), we investigated the impact of *P. multocida* infection on cell apoptosis. Analysis of the expression or cleavage of apoptosis markers (Bcl-2, BAX, Caspase3, Caspase9) as well as annexin V staining revealed that *P. multocida* infection promoted apoptotic cell production, increased the expression of the pro-apoptotic protein BAX, and decreased the expression of the anti-apoptotic protein Bcl-2 (Fig. 6A, B, C, D, E). These findings indicated that *P. multocida* infection indeed induced cell apoptosis. Subsequently, we assessed apoptosis in ABCF2-overexpressed and knockout cells post *P. multocida* infection. The results showed that ABCF2 overexpression elevated the level of cell apoptosis, while ABCF2 knockout significantly reduced the level of cell apoptosis induced by *P. multocida* infection (Fig. 6F, G, H), suggesting that ABCF2 contributed to *P. multocida*-induced cell apoptosis.

**Fig. 6.**
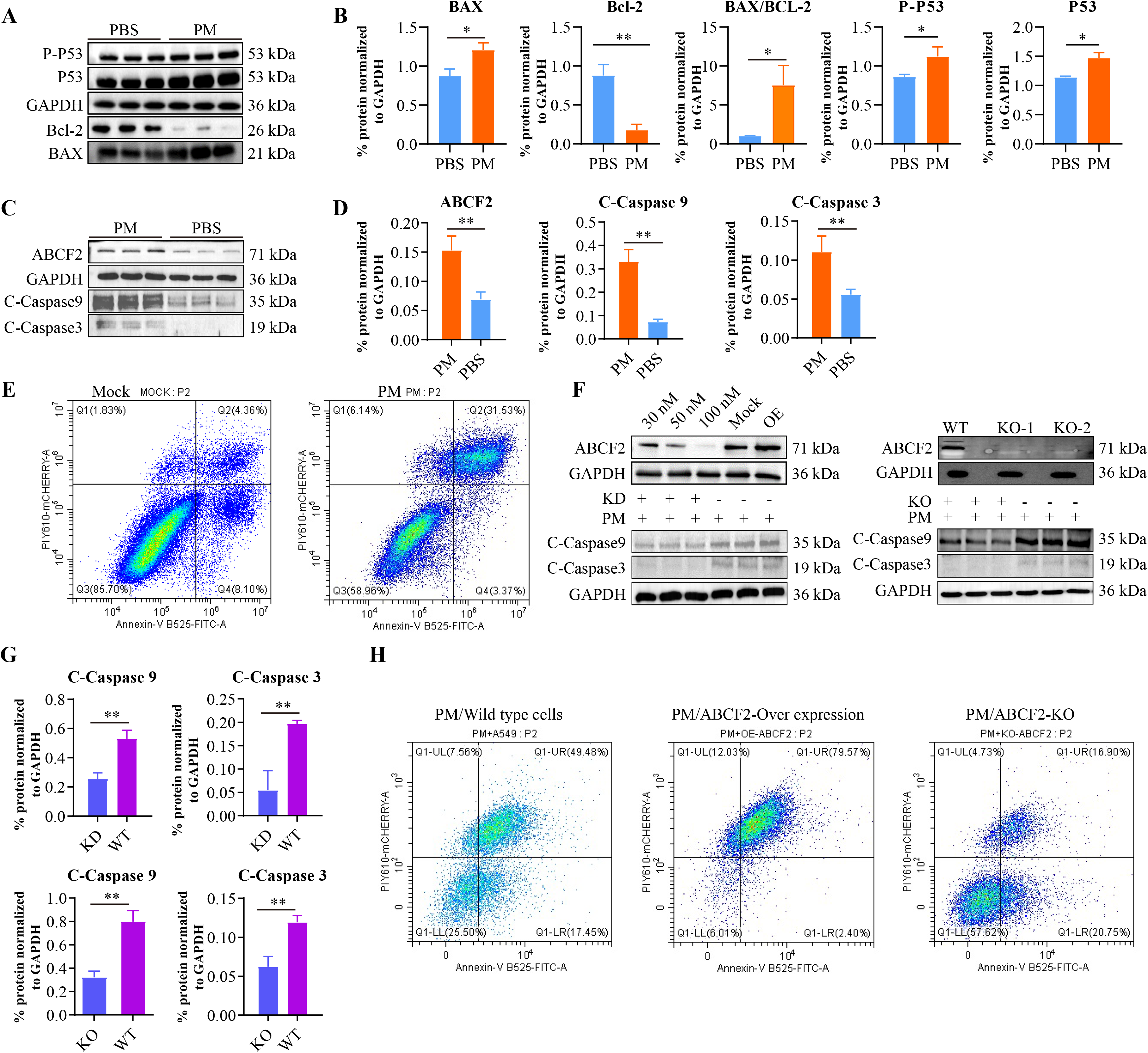
Involvement of ABCF2 in *Pasteurella multocida* induced apoptosis. **(A)** Western-blotting results showing the levels of P53, P53 phosphorylation (P-P53), Bcl-2, and BAX in NPTr cells after *P. multocida* infection. **(B)** The levels of P53, P53 phosphorylation (P-P53), Bcl-2, and BAX in NPTr cells after *P. multocida* infection from western-blotting assays quantified by the image J software. **(C)** Western-blotting results showing the levels of ABCF2, caspase 3 cleavage (C-caspase3) and caspase 9 cleavage (C-caspase9) in NPTr cells after *P. multocida* infection. **(D)** The levels of ABCF2, caspase 3 cleavage (C-caspase3) and caspase 9 cleavage (C-caspase9) in NPTr cells after *P. multocida* infection from western-blotting assays quantified by the image J software. **(E)** Flow cytometry assays detecting apoptotic cells after *P. multocida* infection. **(F)** Western-blotting results showing the levels of caspase 3 cleavage (C-caspase3) and caspase 9 cleavage (C-caspase9) in ABCF2-knocking down (KD), and ABCF2-knocking out (KO) cells after *P. multocida* infection. The knocking down of ABCF2 was achieved using a commercially synthesized siRNA and its efficacy for knocking down at different concentrations (30 nM, 50 nM, 100 nM) was examined using western-blotting. Wild type cells (Mock) and ABCF2-overexpressing cells (OE) were used as controls. The knocking out of ABCF2 was also confirmed using western-blotting. **(G)** The levels of caspase 3 cleavage (C-caspase3) and caspase 9 cleavage (C-caspase9) in ABCF2-knocking down (KD), ABCF2-knocking out (KO), and wild type (WT) cells after *P. multocida* infection from western-blotting assays quantified by the image J software. **(H)** Flow cytometry assays detecting apoptotic cells in ABCF2-overexpressing cells (OE-ABCF2), ABCF2-knocking out (KO-ABCF2), and wild type (A549) cells after *P. multocida* infection. In all column charts, data were presented as mean ± standard deviation (SD). The significance level was set at *P* < 0.05 (*), or *P* < 0.01 (**).

Upon reviewing mass spectrometry data published in a recent article (22), we observed an association between ABCF2 and p53. Given the well-established role of p53-dependent apoptosis, particularly in cancer (23, 24), we further investigated whether p53 was involved in ABCF2-induced cell apoptosis following *P. multocida* infection. Our analysis revealed that *P. multocida* infection significantly increased the expression and phosphorylation of p53 (P-p53) (Fig. 6A, B). Furthermore, overexpression of ABCF2 notably enhanced the expression and phosphorylation of p53 induced by *P. multocida*, while ABCF2 knockdown significantly decreased these levels (Fig. 7A, B, C). Consistently, changes in the levels of apoptosis markers (Bcl-2, BAX) and apoptotic cells aligned with these observations (Fig. 7A, B, C). Additionally, inhibition of p53 in wild-type cells or ABCF2-overexpressed cells also reduced the incidence of apoptosis (Fig. 7D, E), indicating that p53 and its phosphorylation played a role in ABCF2-induced cell apoptosis post *P. multocida* infection. Further analysis demonstrated that ABCF2 could interact with p53 (Fig. 7F, G).

**Fig. 7.**
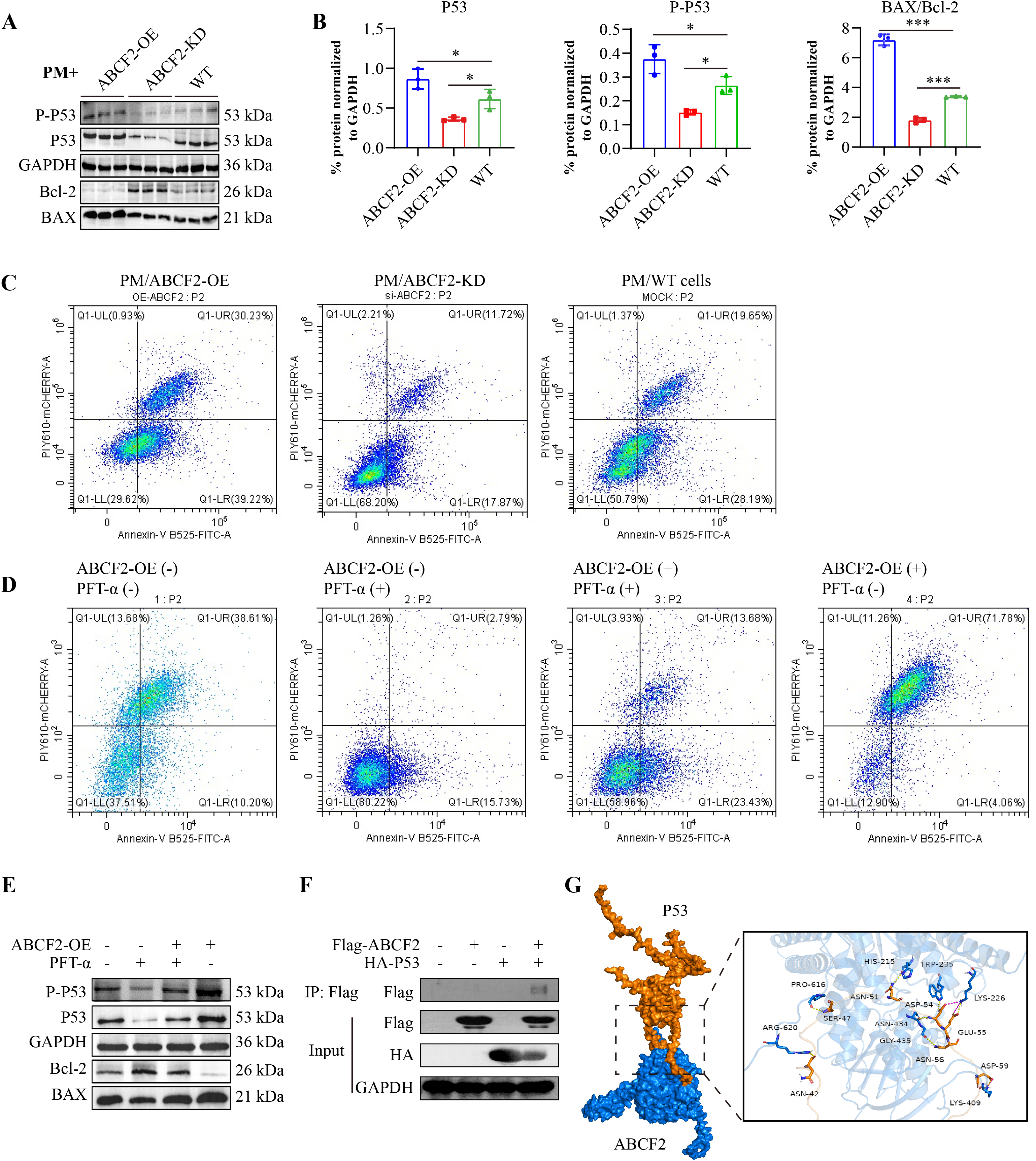
Involvement of ABCF2 in *Pasteurella multocida* induced p53-dependent apoptosis. **(A)** Western-blotting results showing the levels of P53, P53 phosphorylation (P-P53), Bcl-2, and BAX in ABCF2-overexpressing cells (ABCF2-OE), ABCF2-knocking down (ABCF2-KD), and wild type (WT) cells after *P. multocida* infection. **(B)** The levels of P53, P53 phosphorylation (P-P53), Bcl-2, and BAX in ABCF2-overexpressing cells (ABCF2-OE), ABCF2-knocking down (ABCF2-KD), and wild type (WT) cells after *P. multocida* infection from western-blotting assays quantified by the image J software. **(C)** Flow cytometry assays detecting apoptotic cells in ABCF2-overexpressing cells (ABCF2-OE), ABCF2-knocking down (ABCF2-KD), and wild type (WT) after *P. multocida* infection. **(D)** Flow cytometry assays detecting apoptotic cells in ABCF2-overexpressing cells (ABCF2-OE(+)) and wild type (ABCF2-OE(-)) treated with (+) or without (-) p53 inhibitor pifithrin-α (PFT-α) after *P. multocida* infection. **(E)** Western-blotting results showing the levels of P53, P53 phosphorylation (P-P53), Bcl-2, and BAX in ABCF2-overexpressing cells (ABCF2-OE(+)) and wild type (ABCF2-OE(-)) treated with (+) or without (-) p53 inhibitor pifithrin-α (PFT-α) after *P. multocida* infection. **(F)** Validation of the interactions between ABCF2 and P53 by co-immunoprecipitation (Co-IP) assays. **(G)** Validation of the interactions between ABCF2 and P53 by molecular docking analysis. In all column charts, data were presented as mean ± standard deviation (SD). The significance level was set at *P* < 0.05 (*), or *P* < 0.001 (***).

## Discussion

As a long-recognized zoonotic pathogen (confirmed in 1930), *P. multocida* has been responsible for an increasing number of human infections in recent years (25, 26). However, our understanding of the pathogenesis of *P. multocida* remains limited. Numerous review articles and genome-based studies have highlighted various virulence factors that potentially contribute to the pathogenesis of *P. multocida*, with bacterial adherence-associated proteins being a significant focus (5, 7, 13, 14, 27). While the potential involvement of PlpE, PtfA, and/or Hsf-2 in *P. multocida* adhering to host cells has been suggested in several articles (7, 13, 27), their roles as adherence-contributing proteins in *P. multocida* have not been experimentally confirmed. In this study, we experimentally validated the adhesive functions of these three proteins. It is unsurprising to confirm the adhesive roles of PtfA and Hsf-2, considering they are predicted to be type 4 fimbriae and surface fibrils, respectively (27, 28). Both type 4 fimbriae and surface fibrils are recognized as crucial adhesins in various bacterial species (29, 30). Of particular significance is the finding that PlpE also plays a critical role in adhesion of *P. multocida* to host cells in this study. PlpE was initially identified in *Mannheimia haemolytica* (formerly *Pasteurella haemolytica*), a bacterial species closely related to *P. multocida* (31). The homolog of this protein was subsequently discovered in the outer membrane of *P. multocida*, categorizing it as a type of *P. multocida* outer membrane protein (14). Despite the detailed function of this protein not being fully elucidated, several studies have indicated that recombinantly expressed PlpE exhibits significant immunogenic effects (32, 33). Our study findings suggest that PlpE may also function as a bacterial adherence-related protein in *P. multocida*.

In this study, we identified ABCF2 as a host protein that plays roles in the adherence and pathogenesis of *P. multocida*. ABCF2 belongs to the superfamily of ATP-binding cassette (ABC) transporters, and its functions have been extensively investigated in the field of oncology (18, 34, 35). Prior to this report, only one study had reported the involvement of ABCF2 in bacterial pathogenesis (19). In that study, ABCF2 was identified as a target of EspF linked to apoptosis during infections with Enteropathogenic *E. coli* (EPEC) (19). However, unlike the study that observed an anti-apoptotic and protective role against various apoptotic stimuli for ABCF2 during EPEC infection (19), we demonstrated that ABCF2 had a pro-apoptotic role during *P. multocida* infection. Furthermore, we found a positive correlation between ABCF2 and PlpE, as evidenced by increased levels of ABCF2 detected in cells infected with *P. multocida* wild-type cells and *plpE*-complementary strains (CPlpE) compared to those in cells infected with ΔPlpE. This differs from the EPEC study, which found a negative correlation between ABCF2 and EspF, as the *espF* mutant EPEC strain induced increased levels of ABCF2 in cells compared to the wild-type strains and complemented strains (19). These differing observations suggest that ABCF2 may have diverse associations with apoptosis during bacterial infections.

Our analysis revealed that *P. multocida* infection upregulates the level of ABCF2 by activating the NF-κB signaling and p38 MAPK signaling pathways. The regulation of ABCF2 by the NF-κB signaling has been reported in LPS-induced ulcerative colitis (17). Being a pathogen commonly associated with bacterial pneumonia and other inflammatory disorders, *P. multocida* infection is consistently accompanied by the activation of NF-κB signaling, a well-known proinflammatory pathway (36, 37). While the regulation of ABCF2 by the p38 MAPK signaling has not been previously reported in bacterial infections, earlier studies have documented the activation of the p38 MAPK signaling pathway induced by *P. multocida* infection (38, 39). Moreover, the role of the p38 MAPK signaling pathway in modulating the NF-κB signaling was also confirmed in this study, aligning well with findings from other research (21). Thus, the regulation of ABCF2 by the p38 MAPK signaling pathway could potentially occur through the p38 MAPK/NF-κB axis following *P. multocida* infection, with the increased levels of ABCF2 contributing to the adhesion and invasion of host cells by *P. multocida*.

Recently, *P. multocida* infection has been found to induce apoptosis (40, 41). In this study, we demonstrated that ABCF2 is associated with p53-dependent apoptosis induced by *P. multocida* infection. Notably, p53-dependent apoptosis has also been observed in infections caused by various other bacterial species, including *Haemophilus parasuis*, a species closely related to *P. multocida* (42). However, the connection between ABCF2 and p53-dependent apoptosis in bacterial infections has not been reported previously. In numerous non-infectious disease models, p53 has been shown to be activated by inflammatory factors and/or reactive oxygen species (ROS) (43, 44), leading to its upregulation at sites of inflammation (45). Given that *P. multocida* infection typically triggers inflammation in multiple organs such as the lungs and brain, and can also induce ROS production in the lungs (8, 36), p53-dependent apoptosis may play a significant role in *P. multocida* pathogenesis.

In conclusion, we identified ABCF2 as a critical host protein that interacts with three adhesive proteins (PlpE, PtfA, Hsf-2) of *P. multocida*. During infection, *P. multocida* increased the level of ABCF2 by activating the p38 MAPK signaling and NF-κB signaling pathways. The upregulation of ABCF2 contributed to *P. multocida* adhering to and invading host cells. Furthermore, the increased levels of ABCF2 were associated with p53-dependent apoptosis following *P. multocida* infection. The data presented in this study not only enhance our understanding of the pathogenesis of the zoonotic pathogen *P. multocida* but also underscore the involvement of ABCF2 and ABCF2-induced p53-dependent apoptosis in bacterial infections.

## Materials and Methods

### Bacterial strains, cell lines and culture conditions

*P. multocida* strain HuN001 (GenBank CP073238) from a pneumonia patient’s sputum (46) was used alongside strains HB03 (CP003328), HN05 (PPVF01000051), HN07 (CP007040) (47), and *B. bronchiseptica* QH0814 (48), *K. pneumoniae* HB-1, *E. coli* DH5α, and DL21. These strains were sourced from clinical pig (HN05, HN07, QH0814, HB-1) or human samples (HuN001), or commercially (DH5α and DL21). *P. multocida* and *B. bronchiseptica* were cultivated on tryptic soy agar (TSA) or in tryptic soy broth (TSB) (Becton, Dickinson and Co., Sparks, USA) with 5% newborn calf serum (Tianhang, Hangzhou, China). *K. pneumoniae* and *E. coli* strains were grown on Luria-Bertani (LB) agar or in LB broth (Thermo Fisher Scientific, Waltham, USA). NPTr cells, A549 cells, and HEK 293T cells were maintained as previously described (49).

### Construction of bacterial gene deletion and complementary strains

The *plpE* gene was knocked out in *P. multocida* strain HuN001 following a known procedure (50). Upstream and downstream *plpE* fragments were PCR-amplified using specific primers (Table S2), cloned into plasmid pSHK5(TS)-NgAgo to form pSHK-ΔPlpE, and transformed into HuN001 cells via electroporation (2500V, 25μF, 600Ω). Cultures were grown in TSB with 10% calf serum at 28°C for 3 hours, then plated on TSA with kanamycin (50 μg/ml) and serum (5%) for selection. PCR confirmed initial gene replacement with primers PlpE-oL/PlpE-oL (Table S2), followed by overnight culture on TSA at 42°C. Deletion was confirmed via PCR using primers PlpE-iL/iR (Table S2) and Sanger sequencing. For complementation, the full *plpE* sequence was PCR-amplified using primers C-PlpE-F/C-PlpE-F (Table S2), inserted into pPBA1101, and transformed into HuN001-ΔPlpE cells through electroporation.

### TurboID screening

To identify host proteins interacting with PlpE, PtfA, and Hsf-2 from *P. multocida*, their nucleotide sequences were PCR-amplified with specific primers (Table S2). The turboID sequence was also amplified from pET28a-ompP2-turboID and linked to *plpE*/*ptfA*/*hsf-2* fragments via overlap PCR, then cloned into pET-28a(+) to create recombinant plasmids. *E. coli* BL21 cells were transformed with these plasmids and induced to express the proteins using IPTG (0.8mM, Sigma, St. Louis, USA). The recombinant proteins were purified using HisTrap HP columns (GE HealthCare, Chicago, USA). Next, purified proteins (His-TurboID-ptfA, His-TurboID-hsf-2, His-TurboID-plpE, and/or His-TurboID; 3 μg) were incubated with NPTr cells on ice for 1 hour, followed by biotin (50 μM, Sigma, St. Louis, USA) addition and incubation at 37°C for 15 minutes. After cell lysis using a Cell lysis buffer for Western and IP containing a protease inhibitor cocktail (Beyotime, Shanghai, China), the lysate was centrifuged, and supernatants were incubated with streptavidin magnetic beads (NEB, Ipswich, USA) overnight at 4 °C. Biotinylated proteins were eluted by boiling the beads in 100 μL SDT buffer (4% SDS, 100 mM Tris-HCL, 1 mM DTT, pH=7.6), and analyzed via mass spectrometry to identify host cell interactors.

### Cell adhesion and invasion assays

The impact of PlpE, PtfA, and Hsf-2 on bacterial adherence and invasion was assessed using cell models. Full-length *plpE* and *ptfA* were PCR-amplified from *P. multocida* HuN001 using specific primers listed in Table S2, while *hsf-2* was from *P. multocida* HB03. These fragments were cloned into pPBA1101 and transformed into *E. coli* DH5α or *P. multocida* HuN001 to create strains carrying these genes. Next, different cell lines (A549, NPTr, 293T) cultured in wells (2.5×10^6^ cells per well) of a 96-well plate (Corning, USA) were infected with the generated bacterial strains at an optimal multiplicity of infection (MOI) of 200. After incubation at 37°C for 1.5 hours, the cells were washed using PBS, and then lysed in 1 mL of sterile distilled water. To quantify the adherent bacteria, the cells were treated with gentamicin (100 μg/mL) and incubated at 37°C for 1 hour before lysis. Serial 10-fold dilutions of the lysed cells were plated on TSA (for *P. multocida*) or LB agar plates (for *E. coli*) for bacterial enumeration. The number of adherent bacterial strains was calculated as the difference between the bacterial counts from the lysed cells without antibiotic treatment and those with antibiotic treatment. These experiments were independently repeated three times.

ABCF2’s role in bacterial adherence was investigated by amplifying *abcf2* sequences from NPTr and A549 cells, cloning them into pCAGGS-N-flag to generate overexpression plasmids pCAGGS-N-flag-ABCF2-P and pCAGGS-N-flag-ABCF2-H. These were transfected into cells with Lipofectamine 2000 (ThermoFisher, Waltham, USA). Cells transfected with the empty vector pCAGGS-N-flag served as controls. A siRNA (Table S2) was also used to downregulate ABCF2. Cells with ABCF2 overexpression or knockdown were infected with bacterial strains (MOI=200) to study adherence and invasion.

### Laser confocal microscopy

NPTr and A549 cells were grown on confocal dishes (Biosharp, Hefei, China) for 24 hours, then fixed with 200 μL of 4% paraformaldehyde for 20 minutes at room temperature, permeabilized with Triton-X100 for 15 minutes, and blocked with 5% BSA in PBS at room temperature for 2 hours. The cells were treated with an ABCF2 antibody (1:300, MyBioSource, San Diego, USA) at 37°C for 90 minutes, followed by a Cy3-conjugated Goat Anti-Rabbit IgG (H+L) antibody (1:300, Proteintech, San Diego, USA) at 37°C for another 90 minutes. After staining with DAPI (Beyotime, Shanghai, China), the cells were imaged using a Zeiss LSM 800 confocal microscope (Zeiss, Oberkochen, Germany) to visualize ABCF2 expression.

### Molecular docking

Protein-protein interactions were explored via molecular docking to pinpoint crucial amino acids. Structures of PlpE, PtfA, Hsf-2, ABCF2, and p53 were modeled using AlphaFold3 and visualized with PyMOL 3.03. During docking, ABCF2 acted as the receptor, while PlpE, PtfA, Hsf-2, or p53 were ligands. Schrödinger software facilitated docking following a standard protocol (51). Truncated proteins were crafted based on essential amino acids from docking results, and mutations were made with the Mut Express II Fast Mutagenesis Kit V2 (Vazyme, Nanjing, China). Expressing these variants in *E. coli* BL21, purification followed using GE Healthcare HisTrap HP columns (GE HealthCare, Chicago, USA) to delve deeper into their interactions and functions

### Co-immunoprecipitation

To investigate ABCF2 interactions with *P. multocida* proteins and mutants, pCAGGS-N-flag-ABCF2-P and pCAGGS-N-flag were transfected into HEK293T cells using Lipofectamine 2000. After a 24-hour incubation at 37 °C, cells lysis was done with Cell lysis buffer for Western and IP (Beyotime, Shanghai, China), and proteins (2 mg) were incubated with His-tagged proteins (PlpE, PtfA, Hsf-2, mutants, 2.5 μg) for 2 hours at 4°C (step 1). Concurrently, a commercially purchased DYKDDDK-tagged mouse monoclonal antibody (2.5 μg, Proteintech, San Diego, USA) was incubated with rProtein A/G Magarose Beads (2.5 μL, Smart-Lifesciences, Changzhou, China) for 4 hours at room temperature to obtain DYKDDDDK-tagged beads. Subsequently, the protein complex from step 1 was incubated with the DYKDDDDK-tagged beads overnight at 4°C. The beads were then incubated in 3×Flag Peptide (Beyotime, Shanghai, China) for 2 hours at 4°C to elute the proteins, which were later separated by SDS-PAGE. Proteins were transferred onto a PVDF membrane (Bio-Rad, Hercules, USA) and blocked in PBST containing 5% BSA for 3 hours at room temperature. The blots were probed with the DYKDDDK-tagged mouse monoclonal antibody (1:5000, Proteintech, San Diego, USA), His-tag polyclonal antibody (1:5000, Proteintech, San Diego, USA), and/or GAPDH polyclonal antibody (1:5000, Proteintech, San Diego, USA) overnight at 4°C. After washing using PBST, the blots were incubated with HRP-conjugated goat anti-mouse IgG(H+L) (1:5000, Proteintech, San Diego, USA) for 1 hour, followed by being visualized using BeyoECL Star (catalog no. SA00001-1, Beyotime, Shanghai, China).

### Bio-layer interferometry

Bio-layer interferometry (BLI) assays were also conducted to investigate protein interactions. His-tagged proteins (PlpE/PtfA/Hsf-2/truncated/mutated, 50 μg/ml) were immobilized on an anti-His biosensor (ForteBio, Menlo Park, USA). ABCF2 at various concentrations (18.75 nM to 600 nM) was applied, and interactions were analyzed using an Octet RED96 BLI system (ForteBio, Menlo Park, USA).

### CRISPR editing

To knock out ABCF2 in A549 cells, sgRNAs ("aggctgccaaagctcgacag" or "agaagcagacaatcaccaag") were designed and delivered via pSpCas9(BB)-2A-EGFP(Habcf2) plasmid using Lipofectamine 2000. After sorting with a BD FACSAria™ Fusion Flow Cytometer (BD Biosciences, Franklin Lakes, USA), ABCF2-KO cells were obtained from green fluorescence-sorted populations. Confirmation was done via qPCR and western blotting.

### Double luciferase reporting experiment

A luciferase reporter assay studied ATF2 regulation on ABCF2. STAT1 and ATF2 sequences were PCR-amplified from NPTr cell cDNAs using specific primers in Table S2 and cloned into pcAGGS-N-flag. The ABCF2 promoter region was also amplified and inserted into pGL3-Basic. These constructs were transfected into HEK293T cells with Lipofectamine 2000 for analysis using a Dual Luciferase Reporter Assay Kit (Vazyme, Nanjing, China).

### Western blotting

Prior to experiments, cells underwent various treatments: (1) ABCF2 knockdown/knockout verification using siRNA or CRISPR; (2) monolayers of NPTr or A549 cells were inoculated with *P. multocida* HuN001, ΔPlpE, or CPlpE at a MOI = 200 for 12 h to examine the expression of ABCF2; (3) NF-κB signaling after *P. multocida* inoculation in NPTr cells by inoculating NPTr monolayers with *P. multocida* HuN001 (MOI = 200) for 1 h, 2 h, 6h, and 12 h to examine P-P65; (4) NF-κB signaling impact on ABCF2 expression post-*P. multocida* infection by treating NPTr monolayers using an IκB/IKK inhibitor BAY11-7082 (5 μM, Beyotime, China) for 24 h, followed by being inoculated with *P. multocida* HuN001 (MOI = 200) for additional 12 h to examine P-P65 and expression of ABCF2; (5) *P. multocida* effect on p38 MAPK signaling in NPTr cells by inoculating cell monolayers with *P. multocida* HuN001 at a MOI = 200 for 0 h, 4 h, 8h, 12 h, 16 h, and 20 h to examine P-P38; (6) p38 MAPK signaling impact on ABCF2 expression post-*P. multocida* infection by treating NPTr monolayers using an p38 inhibitor BIRB796 (40 μM, Selleck, China) for 12 h, followed by being inoculated with *P. multocida* HuN001 (MOI = 200) for additional 12 h to examine P-P38 and expression of ABCF2. The level of P-P65 was also examined to investigate the 38 MAPK signaling on the NF-κB signaling; (7) ATF2’s regulation on ABCF2 post-*P. multocida* infection by transfecting NPTr cells with pcAGGS-N-flag-ATF2 or pcAGGS-N-flag were inoculated with *P. multocida* HuN001 (MOI = 200) for 12 h to examine the expression of ABCF2; (8) ABCF2’s role in p53-dependent apoptotic signaling following *P. multocida* exposure by inoculating monolayers of wild type cells, ABCF2-KD cells, ABCF2-KO cells, and ABCF2-overexpressed cells with *P. multocida* HuN001 for 12 h to examine the level of p53, Bcl-2, BAX, and the cleavages of Caspase3 and Caspase9, respectively.

Subsequently, cells were lysed using Cell lysis buffer for Western and IP with protease inhibitors, centrifuged at 12000 rpm for 10 min, and subjected to SDS-PAGE. Proteins were transferred to PVDF membranes (Bio-Rad, Hercules, USA), blocked in PBST (containing 5%BSA) for 3 h at room temperature, and probed with specific antibodies overnight at 4 °C to examine the expression of ABCF2 (ABCF2 polyclonal antibody [1:200, MyBioSource, San Diego, USA]), Flag-ATF2 (DYKDDDDK tag polyclonal antibody [1:5,000, Proteintech, San Diego, USA]), p53 (p53 monoclonal antibody [1:10000, Proteintech, San Diego, USA]), P-p53 (phospho-p53 (Ser15) monoclonal antibody [1:2000, Proteintech, San Diego, USA]), Bcl-2 (Bcl-2 polyclonal antibody [1:2000, Proteintech, San Diego, USA]), BAX (BAX monoclonal antibody [1:10000, Proteintech, San Diego, USA]), Caspase3 (Caspase 3/p17/p19 polyclonal antibody [Proteintech, San Diego, USA]), Caspase9 (Caspase 9/p35/p10 polyclonal antibody [Proteintech, San Diego, USA]), and/or GAPDH (GAPDH polyclonal antibody [1:5000, Proteintech, San Diego, USA]). After washing using PBST, the proteins were incubated with HRP-conjugated goat anti-mouse IgG(H+L) (1:5000, Proteintech, San Diego, USA) or HRP-conjugated goat anti-rabbit IgG(H+L) (1:5000, Proteintech, San Diego, USA) for 1 h. Blots were finally visualized with ECL, and quantified using ImageJ (v1.8.0). Immunoreactivity was normalized to loading controls for analysis.

### Flow cytometry

Flow cytometry assays were conducted to investigate the role of ABCF2 in p53-dependent apoptotic signaling post *P. multocida* infection. Wild type cells, ABCF2-KD cells, ABCF2-KO cells, and ABCF2-overexpressing cells were seeded into wells (1 × 10^5^ cells per well) of a 6-well plate (Corning, Corning, USA) and cultured for 12 hours. Wild type cells and/or ABCF2-overexpressing cells were treated with or without the p53 inhibitor pifithrin-α (40 μM, Beyotime, Shanghai, China). Cells were then inoculated with *P. multocida* HuN001 (MOI = 200) for 12 hours. Apoptotic cells were stained using an Annexin V-FITC Apoptosis Detection Kit (Beyotime, Shanghai, China) and analyzed using a CytoFLEX Flow Cytometer (Beckman, Brea, USA).

### qPCR

Primers used for qPCR are listed in Table S2 in supplementary materials. Total RNAs were extracted from bacterial strains at mid-log phase or cells using the TRIzol reagent protocol (Vazyme, Nanjing, China). RNAs were then synthesized to cDNAs using a PrimeScript RT reagent Kit with gDNA Eraser (TaKaRa, Beijing, China) which were used as the templates to examine the transcriptional levels of target genes (*plpE*, *ATF2*, *stat1*, *abcf2*) by PCR with primers and probes listed in Table S2. Bacterial 16sRNA was used as a reference gene for *plpE*, while GAPDH was used as a reference gene for *ATF2*, *stat1*, and *abcf2*.

### Mouse experiments

To evaluate PlpE’s impact on *P. multocida* adherence and invasion, a mouse study was conducted. Briefly, 4∼6-week-old SPF mice (Kunming, female) were randomly divided into three groups and each group contained five mice. Mice in different groups were given an intraperitoneal injection of *P. multocida* wild type strains (100 CFU per mouse), ΔPlpE (100 CFU per mouse) and/or PBS (100 µL per mouse), respectively. After challenging, health conditions of mice were recorded every 2 h, and at 12 h post challenge, three mice from different groups were euthanized to collect the lungs for bacterial counting. The experiment was approved by the Ethic Committee of Huazhong Agricultural University (approval ID number: HZAUMO-2024-0192).

## Statistical analysis

The multiple-t-test strategy in GraphPad Prism 8.0 (GraphPad Software, San Diego, CA, USA) was used to conduct statistical analysis. Data represent mean ± standard deviation (SD). The significance level was set at *P* < 0.05 (*), *P* < 0.01 (**), or *P* < 0.001 (***).

## Acknowledgments

We sincerely acknowledge Prof. Hongbo Zhou at Huazhong Agricultural University for the gift of NPTr cells and A549 cells, Prof. Anding Zhang at Huazhong Agricultural University for the gift of plasmid pSHK5(TS)-NgAgo, Prof. Wentao Li at Huazhong Agricultural University for the gift of plasmid pET28a-ompP2-turboID, and Prof. John D. Boyce at Monash University for the gift of plasmid pPBA1101.

This work was supported in part by Hubei Provincial Natural Science Foundation of China (grant no. 2023AFA094), Fundamental Research Funds for the Central Universities (Project 2662023PY005), Yingzi Tech, and the Huazhong Agricultural University Intelligent Research Institute of Food Health (No. IRIFH202209), the China Agriculture Research System of MOF and MARA (CARS-35), Huazhong Agricultural University and Hubei Hongshan Laboratory Startup Fund. The funders had no role in the study design, data collection and interpretation, or the decision to submit the work for publication.

## SUPPLEMENTAL MATERIAL

**Fig. S1.** Colocalizations between host ABCF2 and *Pasteurella multocida* PlpE.

**Fig. S2.** Adherence of different bacterial species to NPTr cells with ABCF2 knocking down.

**Table S1.** Putative amino acid restudies in PlpE/PtfA/Hsf-2 of *Pasteurella multocida* that are crucial for interacting with ABCF2.

**Table S2.** Primers, probes and oligonucleotides used in this study.

